# MicroRNA-mediated transgenerational plasticity reveals pathways for ocean warming resilience

**DOI:** 10.64898/2026.05.21.727062

**Authors:** Lucrezia C. Bonzi, Jennifer M. Donelson, Rachel K. Shimada, Philip L. Munday, Timothy Ravasi, Celia Schunter

**Affiliations:** The Swire Institute of Marine Science, School of Biological Sciences, The University of Hong Kong, Hong Kong SAR; College of Science and Engineering, James Cook University, Australia; Marine Climate Change Unit, Okinawa Institute of Science and Technology Graduate University, Japan

**Keywords:** Climate change, intergenerational plasticity, non-genetic inheritance, epigenetics, coral reef fish, parental effects, maternal and paternal effects, gamete epigenetic compatibility, multimodal signaling

## Abstract

With anthropogenic climate change driving unprecedented ocean warming and posing a major threat to marine organisms, phenotypic plasticity mediated by epigenetic mechanisms may enable acclimation and persistence across generations. MicroRNAs (miRNAs) are epigenetic regulators that have long been associated with maladaptive stress responses across generations, yet their role in facilitating transgenerational acclimation remains unexplored. To examine miRNAs’ role in transgenerational thermal plasticity, we conducted a fully factorial split-clutch design experiment in the coral reef fish *Acanthochromis polyacanthus.* Parents and offspring were exposed throughout development to control (+0°C) or elevated (+1.5°C) temperatures, and offspring hepatic miRNA and mRNA were profiled. We identified 678 miRNAs in *A. polyacanthus*, including 226 conserved and 452 novel sequences. Direct (developmental) and parental (transgenerational) warming caused similar numbers of miRNAs to alter expression, while mRNA expression was more strongly influenced by parental effects. The two parents had distinct effects on offspring miRNA-mRNA networks, with paternal warming mainly altering offspring metabolism and stress responses, while maternal exposure predominantly influencing immunity and tissue organization. Overall, mismatched parental temperatures created trade-offs in offspring, whereas congruent (biparental) warming led to a unique, non-additive, response with fewer miRNA changes and lack of stress pathway activation. Our findings reveal that miRNAs mediate both developmental and transgenerational plasticity, highlighting the capacity of transgenerational effects to influence climate resilience, while also emphasizing the necessity of accounting for both maternal and paternal contributions when predicting species’ adaptability to ocean warming.

## 1. INTRODUCTION

Due to anthropogenic activities, global environmental conditions are changing at an unprecedented pace, placing organisms under novel and rapidly intensifying selective pressures worldwide (Intergovernmental Panel on Climate Change (IPCC, 2023). Marine populations are increasingly exposed to higher water temperatures, both on average and in fluctuations, with extreme thermal events, such as marine heatwaves, occurring more often and at higher intensity than ever before (Marcos et al., 2025; Oliver et al., 2018), with lasting effects on biodiversity (Hughes et al., 2018; Morosinotto et al., 2025; Smith et al., 2023). To endure under such rapidly changing environments, populations might rely on phenotypic plasticity, which is the ability of organisms to adjust their physiology, morphology, and/or behaviour to maximize their fitness in response to environmental variation. Epigenetic mechanisms - changes in gene regulation that do not alter the DNA sequence - play a key role in acclimation through plasticity, by tuning gene expression quickly and reversibly in response to the environment (Ecker et al., 2018). Remarkably, such epigenetic marks and the associated phenotypic variation can be inherited from one generation to the next via non-genetic inheritance, leading offspring to exhibit phenotypes shaped by their parental environmental experiences, in a process known as transgenerational phenotypic plasticity (TGP; Adrian-Kalchhauser et al., 2020; Liao et al., 2025; McAndry et al., 2025; Perez & Lehner, 2019). However, differences in maternal and paternal non-genetic contributions (Fellous et al., 2022; Meuthen et al., 2024), combined with sexual conflict (Bonduriansky, 2021) and epigenetic incompatibilities arising from asynchronous gametogenesis (Putnam, 2021), might affect how maternal and paternal cues are integrated to determine offspring fitness. In a fish population exposed to an acute marine heatwave, for example, maternal and paternal germlines could be exposed to thermal stress at different stages of gametogenesis, each with distinct windows of epigenetic sensitivity, and thus carry forward different thermal “memories” into the gametes. Offspring must then reconcile these potentially conflicting parental signals to produce a phenotype that is ideally well suited to their own experienced environment. Nevertheless, the relative contribution of each parent to phenotype formation during TGP remains poorly resolved. Addressing this question has significant ecological implications: if TGP is adaptive, it could provide populations and species with time for selection and evolutionary adaptation to occur (Diamond & Martin, 2021; O’Dea et al., 2016), allowing marine populations to persist under prolonged ocean warming conditions (Donelan et al., 2020; Donelson et al., 2018; Munday et al., 2013).

The major epigenetic factors involved in phenotypic plasticity and non-genetic inheritance are DNA methylation, chromatin structure modifications, and several classes of RNAs (Bohacek & Mansuy, 2015; Kelly et al., 2012). MicroRNAs (miRNAs) are short (∼22 nucleotides), single-stranded non-coding RNAs that act as widespread post-transcriptional regulators of gene expression. By binding the complementary sites in the 3’ untranslated regions (UTRs) of messenger RNAs (mRNAs), mature miRNAs typically direct post-transcriptional repression of such transcripts, eventually resulting in gene silencing and reduced protein output (Bartel, 2018; Shang et al., 2023). A single miRNA can target hundreds of different transcripts (Bartel, 2009), and, in humans, more than 60% of protein-coding genes are miRNA targets (Friedman et al., 2009), resulting in miRNAs being involved in basically all cellular functions. Additionally, miRNA expression can be induced by environmental stress, and interact with other epigenetic systems (for example, by influencing transcription factors or chromatin regulators), amplifying and stabilizing regulatory changes (Bartel, 2018; Cecere, 2021; Mendell & Olson, 2012; Yao et al., 2019). In teleosts, miRNAs have roles not only in inherent mechanisms such as development, regeneration, reproduction, metabolism and immunity (Bizuayehu & Babiak, 2014; Zhou et al., 2021; Zhou et al., 2018), but also in the responses to many environmental stressors (Bizuayehu & Babiak, 2014; Raza et al., 2022), including temperature. Studies on zebrafish (*Danio rerio*) and other commercially valuable species have shown that both acute (e.g. Huang et al., 2018; Liu et al., 2024; Ma et al., 2019; Qiang et al., 2017; Vasadia et al., 2019; Xie et al., 2024; Zhao et al., 2024; Zhou et al., 2021) and chronic (Jiang et al., 2024; Sun et al., 2019) exposures to warming elicit dysregulation of fish miRNAome in a variety of tissues, targeting genes involved in energy metabolism, cell fate regulation, as well as inflammatory and immune responses. Because most studies have focused on short-term (hours to days) warming exposures in adult fish, miRNAs involvement is better understood in acute stress responses than in plastic processes underlying longer-term acclimation (Angilletta, 2009), which are key to climate change resilience. Interestingly, when fish were exposed for longer periods, especially during development (i.e. developmental plasticity), heat exposure left long-lasting miRNA signatures, for example in zebrafish (Gavarikar & Craig, 2025; Johnston et al., 2009; van Gelderen et al., 2022), European sea bass *Dicentrarchus labrax* (van Gelderen et al., 2025), as well as Atlantic cod *Gadus morhua* (Bizuayehu et al., 2015), possibly indicative of persistent phenotypic regulation.

Beyond their roles in acute stress responses and developmental plasticity, the emerging contribution of miRNAs in transgenerational phenotypic plasticity may be even more critical for long-term population persistence in changing environments. Indeed, both small non-coding RNA and their amplification agents (proteins or corresponding gene regulatory states) can be transmitted from parents to offspring (Alcazar et al., 2008; Rechavi et al., 2011; Rodgers et al., 2015) and studies in rodents demonstrate that sperm miRNAs mediate paternal transmission of diet-induced metabolic disorders and stress-related phenotypes (Chen et al., 2016; Dupont et al., 2019; Gapp et al., 2014; Rodgers et al., 2015; Wang et al., 2021), while maternal stress alters miRNA profiles in subsequent generations (Yao et al., 2014). Similar mechanisms might operate in fish, with paternal stress affecting offspring phenotype in zebrafish possibly via modifications in sperm miRNAs (Ord et al., 2020). Furthermore, F0 environmental experiences can leave persistent miRNA signatures in adult F1 and F2 generations (Luu et al., 2021), with both distinct and overlapping maternal and paternal influences reflected in embryonic miRNA expression patterns (Heinrichs-Caldas et al., 2023). While miRNA changes have been traditionally linked to stress states (but see Yin et al., 2025), and could therefore be maladaptive for offspring, their potential role in facilitating transgenerational adaptive responses to environmental change remains largely unexplored.

Here, we aim to elucidate the possible involvement of miRNAs in transgenerational phenotypic plasticity in the context of ocean warming acclimation in the spiny chromis damselfish *Acanthochromis polyacanthus*. In this species, developmental exposure to elevated temperatures leads to partial acclimation to warming (Donelson et al., 2011), while biparental developmental thermal exposure induces full compensation of the metabolic challenges of warming in the offspring (Bernal et al., 2018; Donelson et al., 2012). Such acclimation is mainly achieved through molecular changes in energy metabolism, immune system and transcriptional regulation (Bernal et al., 2022; Bonzi et al., 2024; Ryu et al., 2018; Veilleux et al., 2015). While the phenotypic scope of *A. polyacanthus* ocean warming acclimation potential have been previously investigated, the epigenetic mechanisms that underlie the transgenerational thermal acclimation remain unresolved. Notably, prior work reporting differential DNA methylation associated with thermal exposure (Ryu et al., 2018) is difficult to interpret for true transgenerational inheritance because eggs were exposed to parental conditions during gametogenesis and embryogenesis. Building on our fully factorial, split-clutch design that separated biparental, paternal-only, maternal-only and developmental-only exposures to control or elevated (+1.5°C) temperature while measuring offspring phenotypes and hepatic gene expression (Bonzi et al., 2025; Spinks et al., 2021, 2022), we here profile hepatic miRNA expression across those exposure groups (Fig. 1). We aim to test two main hypotheses. First, we hypothesize that both developmental and parental (transgenerational; *sensu* McAndry et al., 2025) exposure to warming elicit changes in the miRNA profiles of fish livers. If this is the case, we would also expect to identify sex-specific paternal and maternal signatures on offspring miRNA expression and their interactive, additive, or redundant nature in influencing offspring phenotypes. Second, we hypothesize that the genes targeted by these temperature-responsive miRNAs are involved in acclimation processes to warming (e.g. energy metabolism, immunity, transcriptional regulation). Ultimately, this study aims to uncover if and how miRNAs mediate transgenerational responses to warming and how mismatched maternal and paternal cues affect gamete epigenetic compatibility, enhancing our knowledge on the mechanisms of adaptive acclimatory capacities to global change.

**Figure 1.**
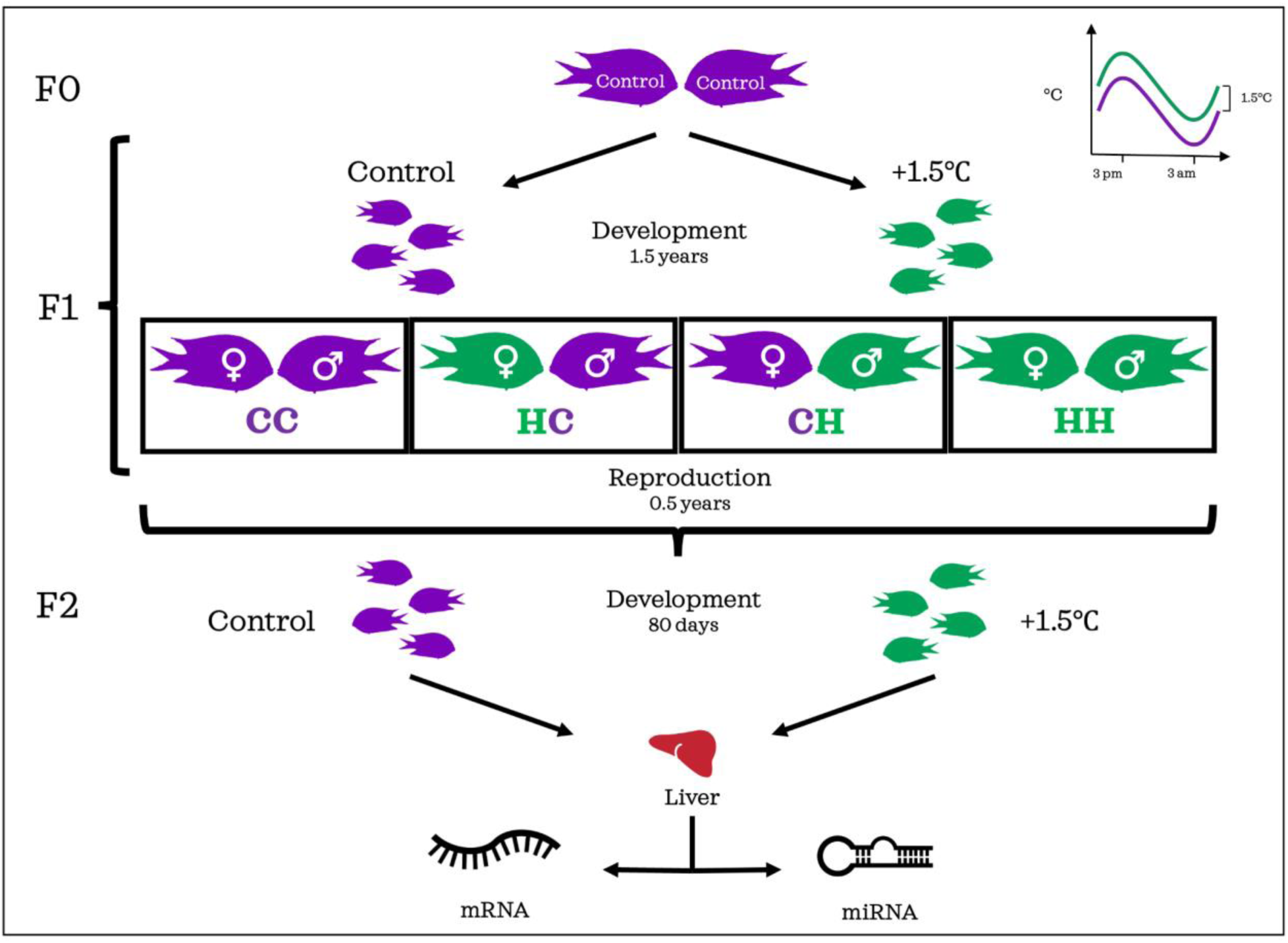
Experimental design. F1 *A. polyacanthus* from wild-caught F0 developed either at present-day control temperature (control - purple) or elevated temperature (+1.5°C - green), with seasonal and daily fluctuations. At 1.5 years of age, F1 individuals were paired in reciprocal sex crosses of the two thermal treatments and kept at control temperature until reproduction. F2 siblings from the four parental thermal combinations were split after hatching into control or elevated temperatures, where they developed for 80 days. F2 livers (n = 8-12 fish per treatment) were collected for total RNA extraction, and mRNA and miRNA sequencing. Treatment code: first letter = thermal treatment of the father; second letter = thermal treatment of the mother; C = control temperature; H = +1.5°C elevated temperature

## 2. METHODS

### 2.1 Experimental design

To explore the putative role of miRNAs in transgenerational plasticity in response to ocean warming and understand the individual contributions of fathers and mothers, we exposed *Acanthochromis polyacanthus* to increased water temperatures over two generations, as detailed in Spinks et al. (2021, 2022). Six wild adult pairs (F0 generation) were collected from the Palm Island region (18°37′ S, 146°30′ E) and nearby Bramble Reef (18°22′ S, 146°40′ E) of the central Great Barrier Reef in 2014/2015. F0 were kept at seasonally cycling, present-day temperatures resembling those of the collection site at the Marine and Aquaculture Research Facility of James Cook University (Townsville, Australia). In the Austral summer of 2016/2017, the F1 was produced, and clutches were split into two different rearing temperature treatments as soon as the larvae hatched, with at least five replicate tanks per thermal treatment. The control temperature treatment simulated seasonal (winter minimum 23.2°C, summer maximum 28.5°C) and diurnal (constant cycling between daily minimum −0.6°C at 3.00 AM and daily maximum +0.6°C at 3:00 PM) cycles for the Palm Islands region. The elevated temperature treatment matched the above cycling, but 1.5°C higher, which is the expected average ocean temperature increase by the end of the century in a low CO_2_ emission scenario (Fox-Kemper et al., 2021). At 1.5 years of age, F1 fish reached sexual maturity and non-sibling breeding pairs were formed so that reciprocal sex crosses of the developmental temperatures were created, resulting in four parental thermal treatment pair combinations: (1) both male and female developed at control temperature (CC), (2) male developed at elevated temperature and female developed at control temperature (HC), (3) male developed at control temperature and female developed at elevated temperature (CH) or (4) both sexes developed at elevated temperature (HH; Suppl. Table S1). At this time, temperature was adjusted to present-day conditions so that spermiogenesis and oocyte maturation occurred for all individuals at control temperature. Breeding occurred in the Austral summer of 2017/2018, and newly hatched F2 generation siblings were split into present-day control or +1.5°C temperature treatments, matching the above-mentioned seasonal and diurnal cycling. For each clutch, two tank replicates per thermal treatment were set up. F2 fish were reared in any of the two temperature treatments until 80 days post-hatching, when they were sexed by urogenital papilla external examination. A minimum of 3 females and 3 males from at least 3 clutches per treatment were then euthanized by cervical dislocation, measured for standard length, weighed, and dissected, for a total of 77 sampled individuals (8-12 samples/treatment; Supp. Table S2). In addition, another 98 F2 individuals and 23 of their parents (F1) were sampled, for a total of 175 biological replicates, to build a more comprehensive miRNA library for *A. polyacanthus*.

### 2.2 RNA extraction and sequencing

Livers were immediately snap frozen in liquid nitrogen and stored at -80°C. We chose to analyse the liver because of the fundamental role of this tissue in metabolism and to allow comparisons with previous works (Bernal et al., 2018; Veilleux et al., 2015). Samples were processed as described in (Bonzi et al., 2024). Briefly, total RNA was extracted using a mirVana miRNA Isolation Kit, following the total RNA isolation protocol, and checked for quality and quantity. Each sample was then split in two aliquots, one used for RNAseq (described in Bonzi et al., 2025), the other for small RNAseq. Small RNA libraries were prepared with the Illumina TruSeq Small RNA Library Preparation Kit and checked for quality and quantity using a Bioanalyzer High Sensitivity DNA assay (Agilent). For a subset of samples, two separate small RNA libraries were produced and individually sequenced to get some technical replicates. 50 bp single-end read sequencing was performed with an Illumina HiSeq 4000 at the King Abdullah University of Science and Technology Bioscience Core Lab, randomly assigning each sample to different lanes to avoid sequencing batch effects.

### 2.3 miRNA prediction and annotation

Clean reads were obtained from raw data using Cutadapt (v4.4; Martin, 2011) and Trimmomatic (v0.39; Bolger et al., 2014) to remove adapters, low-quality reads (Q ≤ 30), reads without 3*’* adapters, polyA reads, as well as reads shorter than 18 nt and longer than 34 nt. Quality control checks were performed pre- and post-filtering with FastQC (v0.11.9; Andrews, 2010) and mirnaQC (Aparicio-Puerta et al., 2020). The obtained high-quality sequences were then aligned using Bowtie (v1.3.1; Langmead et al., 2009) to a library of small non-coding RNAs (rRNAs, tRNAs, piRNAs, snoRNAs and snRNAs) sourced from Rfam (Kalvari et al., 2021), piRNABank (Sai Lakshmi & Agrawal, 2008) and GenBank (Benson et al., 2008). Reads that aligned with the negative reference library were removed, while the remaining sequences were used for miRNA identification with miRDeep2 (v2.0.1.3; Friedländer et al., 2012). Specifically, miRDeep2 mapper.pl module (options: -d -e -i -j -m) was first used to collapse and map all the combined filtered reads from all our samples to *Acanthochromis polyacanthus* reference genome (ENSEMBL ASM210954v1). The resulting alignment file and processed reads were then used as input to miRDeep2.pl wrapper function to detect and predict hairpin-like miRNA precursors in our data. To guide the annotation, we used a set of known mature miRNAs sequences of five teleost species (*Danio rerio*, *Oreochromis niloticus*, *Neolamprologus brichardi*, *Oryzias latipes* and *Astatotilapia burtoni*) retrieved from miRbase (v22.1; Kozomara & Griffiths-Jones, 2014). All miRNAs precursors with a significant randfold P-value and miRDeep2 score above 3 were retained. Next, we used BLAST+ (v2.15.0; Camacho et al., 2009) to annotate miRNAs by blasting mature and star miRNA sequences against FishmiRNA (release Sept2023; Desvignes et al., 2022) and miRbase databases. miRNAs with a hit (evalue < 0.01) were considered conserved, while those with no hit were considered novel. To reduce redundancy, all precursors, as well as mature and star miRNAs were blasted against themselves to find identical or similar sequences (evalue < 0.01 for mature and star miRNAs, evalue < 0.005 for precursors). The resulting list of conserved miRNAs was named according to miRNA nomenclature guidelines (Desvignes et al., 2015), whereas a prefix “novel” was used for novel miRNAs.

### 2.4 miRNA differential expression analysis

A matrix of mature miRNAs raw counts was created with the *quantifier.pl* function of miRDeep2, considering multi-mapping since some miRNAs have more than one precursor. For the subset of samples that were sequenced twice from two different libraries each, counts from the two replicates were averaged and treated as single biological samples. Differentially expressed (DE) miRNAs were then identified with DESeq2 (v1.40.2; Love et al., 2014) in R (v4.3.1; R Core Team, 2023). First, the variance-stabilizing transformed counts were visualized with principal component analyses (PCAs) and heatmaps of the sample-to-sample distances to check for outliers and batch effects. As a result, two outlier samples were excluded from further analyses (Suppl. Table S2). Likelihood ratio tests were then run to choose the best model formula and quantify the overall effects of the paternal, maternal and offspring’s own exposures to warming, as well as the presence of potential interactions between these three main factors. Family line and size (defined as weight divided by standard length) were included in the final model, as well as the interaction term between maternal and paternal treatments, in addition to the three main effects of paternal, maternal and F2 offspring thermal experiences. We removed the interactions between any of the parental treatments and the offspring treatment because they were not significant (i.e. they returned no DE miRNAs, adjusted *p*-value < 0.05). To take into account the existence of the interaction between the paternal and the maternal thermal experiences, we then ran pairwise comparisons (contrasts) between offspring from the four possible parental combinations: both parents raised at control temperature (CC), father at elevated temperature and mother at control (HC), father at control and mother at elevated temperature (CH), and both parents raised at elevated temperature (HH). Additionally, we also tested the comparison between all the offspring raised at elevated (F2 H) vs control temperature (F2 C). Within these pairwise comparisons miRNAs with adjusted *p*-value < 0.05 (Benjamini & Hochberg, 1995) and absolute log_2_ fold-change (LFC) > 0.3 were identified as differentially expressed.

### 2.5 mRNA differential expression analysis

The same 77 samples used for DE miRNA analysis were also used to calculate differentially expressed (DE) mRNAs. First, mRNA-seq raw reads were trimmed for quality and adapter removal with Trimmomatic (v0.39; Bolger et al., 2014) and the clean reads were then mapped to the *A. polyacanthus* reference genome with HiSAT2 (v2.2.1; Kim et al., 2019). Then, we used featureCounts (Liao et al., 2014) from the Subread v2.0.2 package to quantify mRNA counts. Finally, similarly to miRNA data above, DESeq2 was used to detect differentially expressed transcripts across treatments (*p*adj < 0.05, |LFC| > 0.3, baseMean > 10) by running pairwise comparisons between offspring from the four different parental combinations (CC, HC, CH and HH), regardless of their own thermal experience, as well as between offspring raised at elevated *vs* control temperature (F2 H vs C), regardless of their parental thermal experience, followed by LFC shrinkage by apeglm method (Zhu et al., 2019).

### 2.6 miRNA target analysis

Putative target transcripts of the differentially expressed miRNAs were predicted following a stringent pipeline that integrates the results of three in silico prediction tools: Miranda (v3.3a), an algorithm that identifies target genes based on sequence complementarity and free energies of the mRNA:miRNA duplexes (Enright et al., 2003); TargetScan (v6.0), which scores predicted targets based on six different site context features (Garcia et al., 2011); and RNAhybrid (v2.1.2), that calculates the energetically most favourable binding sites between miRNAs and their potential targets (Rehmsmeier et al., 2004). Miranda and TargetScan were run with default parameters on the 14,090 3’ UTR sequences available for the *A. polyacanthus* genome. To increase RNAhybrid accuracy, we first applied the RNAcalibrate tool to estimate the distribution parameters specific to our data and used energy −20 kcal/mol as cut-off. All tools were downloaded and used locally. Since a higher expression level of a miRNA should correspond to a lower expression level of its direct targets, the resulting potential miRNA-transcript targeting pairs successfully predicted by all three software were then filtered based on negative correlations (Spearman’s ρ < -0.3) between expression levels of paired transcripts and DEMs across the same 77 samples used for differential expression analyses. Multiple testing correction was performed for each DEM with more than one predicted target, and only correlations with adjusted *p*-values < 0.05 were retained. Functional enrichment analyses for each pairwise comparison list of validated targets were performed in OmicsBox (v3.4.5; BioBam Bioinformatics, 2019) with Fisher’s exact test (FDR < 0.05) and DAVID Functional Annotation tool (Sherman et al., 2022) was used to summarize gene functions and pathways. DEM–mRNA networks were constructed using Cytoscape (v3.10.3; Shannon et al., 2003).

## 3. RESULTS

### 3.1 Identification of miRNAs in A. polyacanthus

To assemble a miRNA reference for *A. polyacanthus*, we built small RNA libraries from liver tissue of 175 individuals, differently exposed to control (+0°C) or elevated (+1.5°C) water temperatures during post-hatching development. A total of ∼2.5 billion raw reads were generated, which, after adaptors and quality filtering, resulted in an average of 8,159,766 clean reads per sample, with a length of 18-34 nt (Suppl. Table S3), peaking between 21 and 22 nt (Suppl. Fig. S1A). Reads peaking at longer lengths were removed after mapping to a negative list of non-miRNA small RNAs, with only the unaligned reads retained and used for miRNA identification (Suppl. Fig. S1B). A set of 892 miRNA precursors was predicted by miRDeep2. After removing redundancy, 678 unique mature miRNAs were identified, including 226 conserved miRNAs, with orthologous sequences found in other species, and 452 novel to *A. polyacanthus* (Suppl. Table S4).

### 3.2. Transgenerational and developmental thermal effects induce distinct miRNA-mRNA regulatory networks

To identify miRNAs involved in transgenerational plasticity in response to ocean warming, we analysed the differential expression (DE) of liver miRNAs in *A. polyacanthus* offspring from different parental and personal thermal experiences (Fig. 2). Overall, a similar magnitude of responses was elicited by the offspring’s direct experience of warming (28 DE miRNAs) and by the transgenerational one (father: 17 DE miRNAs, mother: 14 DE miRNAs, 8 shared). No interaction between the offspring’s own thermal experience and that of any of its parents was found, but six miRNAs were differentially expressed because of the interaction between the paternal and the maternal exposures to warming (Fig. 2A). Pairwise comparisons between offspring from different parental combinations (HC vs CC, CH vs CC, HH vs CC) and between offspring raised at elevated vs control temperature (F2 H vs C) similarly showed comparable numbers of DE miRNAs due to the personal experience of warming (23 DE miRNAs) and due to the transgenerational one (total of 21 DE miRNAs due to the different parental exposures; Table 1; Suppl. Fig. S2; Suppl. Table S5). On the contrary, the mRNA differential expression analysis revealed a different picture, with larger numbers of mRNAs differentially expressed because of the parental experience compared to the offspring’s direct exposure to warming (Fig. 2C; Suppl. Table S6).

**Figure 2.**
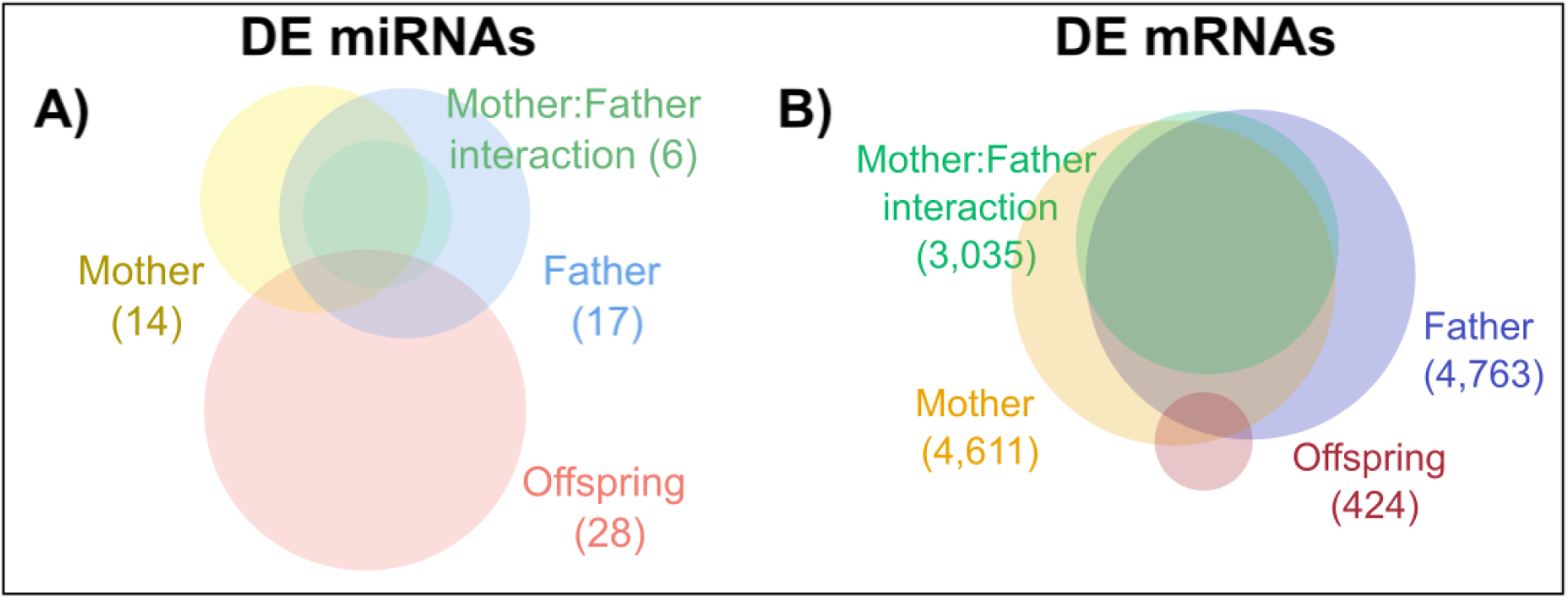
Venn diagrams of the RNA differential expression (DE) in *A. polyacanthus* in response to transgenerational and direct developmental warming. A) DE miRNAs and B) mRNAs attributable to the paternal, maternal and offspring’s own developmental thermal experiences and to the interaction between the paternal and the maternal ones, identified by likelihood ratio tests.

**Table 1.** Results of the pairwise comparisons between offspring from the different parental and personal thermal experiences.

|  | HH vs CC | HC vs CC | CH vs CC | F2 H vs C |
| --- | --- | --- | --- | --- |
| Differentially expressed miRNAs | 5 | 5 | 14 | 23 |
| Differentially expressed mRNAs | 91 | 2,422 | 1,337 | 269 |

To assess whether the differentially expressed miRNAs exerted inhibitory effects on the fish gene expression, we predicted their targets by integrating results from three distinct in silico tools (Suppl. Fig. S3). Subsequently, we evaluated how many of these predicted targets showed a significant negative correlation with the expression levels of their corresponding miRNA in our samples (Suppl. Table S7). This resulted in a total of 656 validated DE miRNA-target mRNA pairs (Fig. 3A, Suppl. Tables S7 & S8). The miRNA apo-miR-101a-3p, DE in offspring from parents where the father alone was exposed to warming (HC) compared to offspring from control (CC) parents, was correlated to the largest number of target mRNAs (167). On the other hand, almost half (49%) DE miRNAs were not found to be significantly correlated to any of the predicted target mRNAs (after multiple testing correction), including apo-miR-459-5p, the only miRNA that was differently expressed in all comparisons. Functional enrichment of the target mRNAs for each pairwise comparison (Fig. 3B) revealed no significant GO terms.

**Figure 3.**
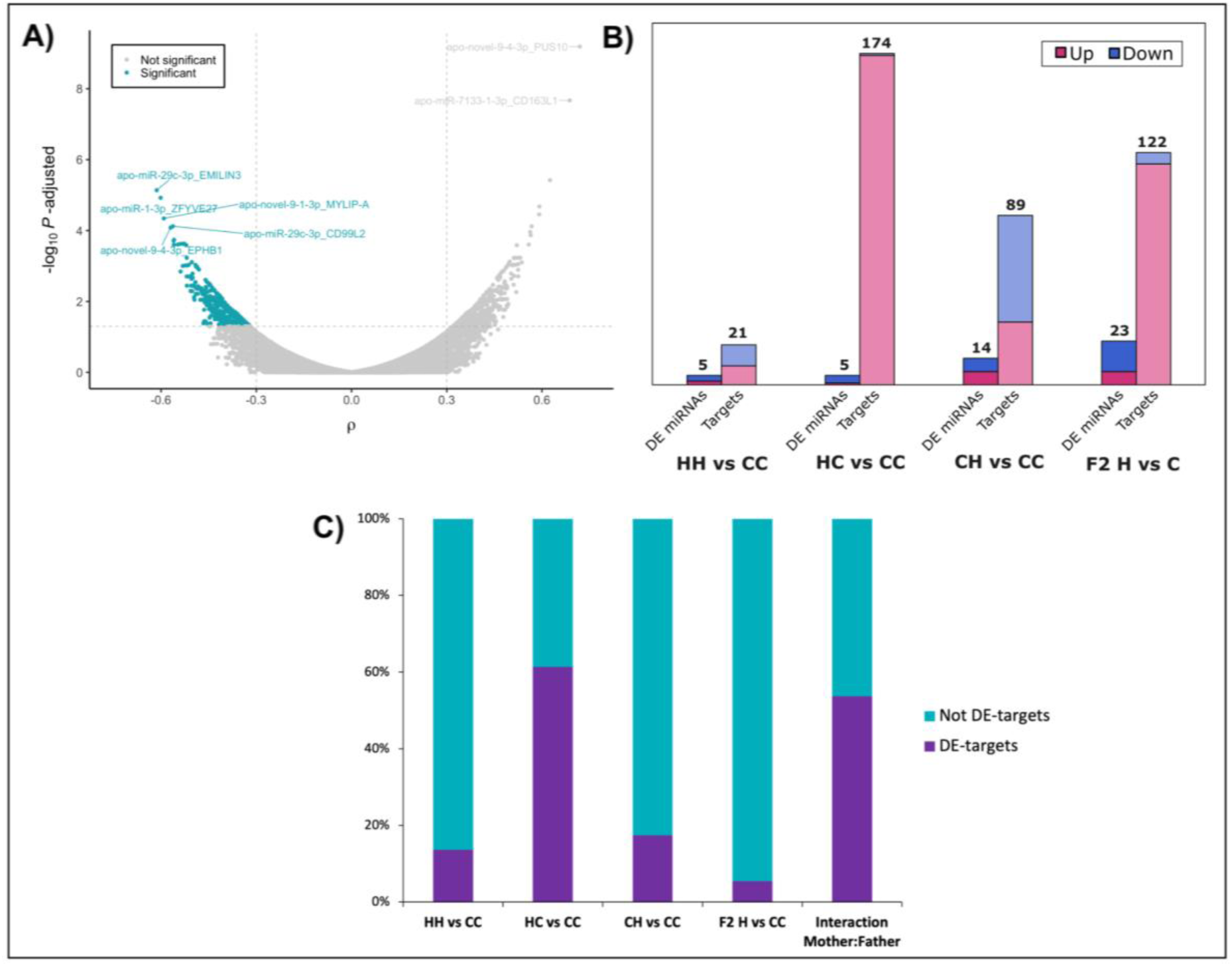
Prediction of high-confidence targets of differentially expressed (DE) miRNAs. A) Predicted DE miRNA-target mRNA pairs filtered by negative correlation (Spearman’s ρ < -0.3, adjusted *P*-value < 0.05) between expression levels of paired mRNAs and miRNAs across all samples, generating 656 validated pairs. B) Up- and down- regulated DE miRNAs and respective target mRNAs per pairwise comparison. C) Proportion of target mRNAs also differentially expressed in each of the pairwise comparisons and due to the interaction between maternal and paternal thermal treatments. The first letter stands for the paternal thermal environment, the second for the maternal one; ‘C’ = control temperature and ‘H’ = +1.5°C.

Finally, we checked if any of the target mRNAs were also significantly differentially expressed in the same samples, and we found the highest proportion of DE target mRNAs in offspring from fathers alone exposed to warming (HC) compared to offspring from control parents (CC), where 109 out of the 176 target mRNAs were differentially expressed (Fig. 3C). Overall, the DE miRNAs due to the paternal exposure to warming regulated 7% of the total DE mRNAs (among the 3’ UTR annotated ones) in the same samples. For the other pairwise comparisons (HH vs CC, CH vs CC, HC vs CC, F2 H vs C) the percentages of the DE mRNAs regulated by DE miRNAs ranged between 0.5-4% (Suppl. Table S9).

#### 3.2.1 Parental effects of warming on offspring miRNA expression

##### 3.2.1.1 Interaction between paternal and maternal thermal exposures

We found that parental thermal histories jointly shaped offspring miRNA expression through interactive effects, with one parent’s contribution modulated by the other’s. Specifically, the expression levels of six miRNAs varied according to distinct maternal-paternal thermal history combinations (Suppl. Fig. S4; Suppl. Table S5). For example, offspring from parents raised at different temperatures (HC and CH) expressed lower levels of certain miRNAs (apo-miR-19b-3p, apo-miR-19da-3p, apo-novel-2-5p and apo-novel-20-5p) compared to those with parents that developed in the same thermal regime (CC and HH; Fig. 4A). Accordingly, their target mRNAs were more highly expressed in offspring from mismatched parents compared to offspring from parents both raised either at control or elevated temperature (Fig. 4B). These target genes are involved in cell proliferation and apoptosis (e.g. *nf2*, *notch2*, *srrt*), stress response (e.g. *arfgap1*, *taok1*, *traf3ip2*), RNA processing, such as mRNA splicing (e.g. *celf1*, *ddx17*, *pnisr*, *rbm25*) and energy metabolism (e.g. *aco2*, *ampd2*, *cpne3*; Suppl. Table S10). Our results therefore indicate that the effect of parental warming exposure on offspring miRNA expression is context-dependent: the thermal history of one parent (mother or father) does not act in isolation but interacts with the other parent’s thermal experience to shape the offspring’s miRNA profile. For this reason, we focused the subsequent analyses on the effects of the four different parental combinations on offspring miRNA transcriptional profiles.

**Figure 4.**
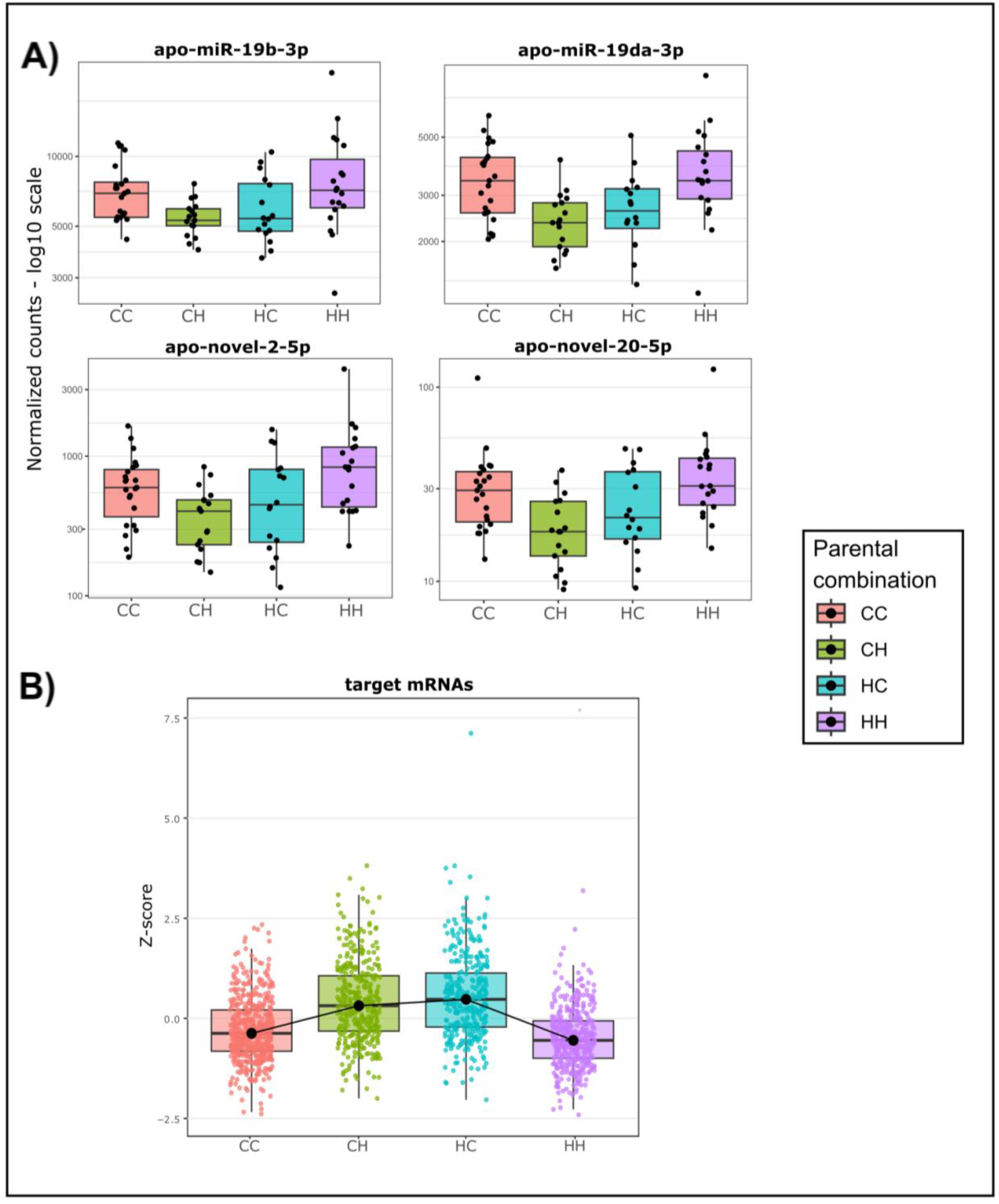
Interactive effects between maternal and paternal thermal histories. Expression profiles of A) a subset of the differentially expressed miRNAs showing interactive effects between maternal and paternal thermal exposures and B) their target mRNAs. In the parental combinations, the first letter stands for the paternal thermal environment, the second for the maternal one; ‘C’ = control temperature and ‘H’ = +1.5°C.

##### 3.2.1.2 Paternal effects of warming (HC vs CC results)

Offspring whose father experienced warming, while the mother control temperature (HC), showed changes in five miRNAs compared to offspring from control parents (CC), regardless of their own developmental temperature (Suppl. Table S5; Suppl. Fig. S5A). One of these, apo-novel-4i-2-3p, was upregulated, and targets the gene *nt5dc2*, which negatively regulates catecholamine synthesis, and was also differentially expressed (Suppl. Table S11). Among the other DE miRNAs, which were expressed instead at lower levels in HC offspring compared to CC ones, apo-miR-101a-3p and apo-miR-375-3p negatively correlated with 173 mRNAs, with 106 of these targets also differentially expressed and upregulated in HC offspring. Many of these mRNAs are involved in metabolism and energy homeostasis (e.g. *acsl6*, *angptl3*, *aqp9*, *frzb/sfrp3*, *nucb2*, *pfkfb1*, *prkab1*, *slc2a8*), as well as transcription and gene expression regulation (e.g. *gatad2b*, *prdm11*, *pura*, *tbp*, *znf23, znf346, znf652*). Other target mRNAs had roles in response to stress, in particular components of the unfolded protein response (e.g. *calr*, *cnpy2*, *edem1*, *hif1an*, *stc2*, *tmx2*, *txndc5*), immune system (e.g. *cybb*, *enpp4*, *leap2*, *marchf8*, *usp12*), vesicle-mediated transport of lipid and proteins (e.g. *arfgap3*, *myo18a*, *rab11fip2*), cytoskeletal regulation, cell proliferation and wound healing (e.g. *acvr1b*, *arhgap35*, *fgf10*, *lox*, *prox1*, *zbtb5*). The target mRNAs that were not differentially expressed played similar roles, particularly in lipid metabolism and energy production, stress and immune response, cell proliferation and differentiation (Suppl. Table S11). Taken together, these coordinated miRNA–mRNA changes indicate that paternal warming affected fundamental physiological processes, with particularly strong impacts on offspring metabolic pathways, transcriptional regulation and stress response systems.

##### 3.2.1.3 Maternal effects of warming (CH vs CC results)

When mothers were exposed to elevated temperature during development, but fathers were not (CH), their offspring differentially regulated 14 miRNAs (7 upregulated and 7 downregulated) compared to offspring from control parents (CC), regardless of the offspring’s own thermal experience (Suppl. Table S5; Suppl. Fig. S5B). Among the upregulated DE-miRNAs, apo-miR-1-3p, apo-miR-27b-1-3p, and apo-novel-37-3p targeted 56 mRNAs, including 10 also differentially expressed and downregulated in CH offspring (Suppl. Table S12). These DE genes play roles in immunity and inflammation (*card8*, *card11, tmed1*), cell junction (*cdh17*) and extracellular matrix organization (*emilin3*), cell growth (*pttg1p*), glycosylation (*dpm1*), endocytic recycling and cholesterol homeostasis (*ehd1*), signalling (*gfra4*), as well as cellular respiration (*nubpl*). The other targeted genes, not differentially expressed, similarly span functions from immune and inflammatory responses (e.g. *cd99l2, havcr1, inpp5d, ninj1*), to cell adhesion and cell proliferation (e.g. *ctnnal1, ddr1, itga5*), stress response (e.g. *dnajb14, lamp3, ppm1l, rbm7*), as well as metabolism and energy regulation (e.g. *akt1*, *galm*, *hmga1*, *slc2a8*) Among the downregulated miRNAs in CH offspring, only apo-miR-19da-3p (also affected by the interaction of maternal and paternal thermal exposures) had mRNAs targets that were significantly negatively correlated with its expression level. Six of the 33 targets were also DE in the same offspring, including genes involved in Golgi enzyme recycling and protein processing (*fam114a1, mansc1*), cell proliferation and apoptosis (*nf2*, *pcdh17*), cytoskeleton and extracellular matrix regulation (*sh3pxd2b*), and innate immunity and inflammatory response (*traf3ip2*). Some of the non-DE target functions were similar, with genes involved in intracellular vesicle trafficking and cytoskeleton regulation (e.g. *agap1*, *iqsec1*, *synrg*), protein processing, and cell fate determination, while other genes were involved in energy metabolism, mRNA splicing (e.g. *celf1, ddx17, rbm25*) and histone acetylation and transcription regulation instead (e.g. *ankrd11*). Overall, these results indicate that maternal thermal exposure induces subtle but significant miRNA-mediated gene regulation in the offspring, mainly influencing immune response, cellular structure, and metabolic pathways, though with less impact than paternal thermal effects.

##### 3.2.1.4 Biparental warming exposure effects (HH vs CC results)

When both parents experienced warming (HH), their offspring differentially expressed five miRNAs compared to offspring from control parents (CC), regardless of the offspring’s own developmental temperature (Suppl. Table S5; Suppl. Fig. S5C). Four of these DE miRNAs were unique to this parental combination: two were downregulated (apo-novel-4c-3p and apo-novel-77-5p), and two upregulated (apo-novel-9-1-3p and apo-novel-9-4-3p). Among the 11 target mRNAs of the upregulated miRNAs in HH offspring, only erb-B2 receptor tyrosine kinase 2 (*erbb2*) was also differentially expressed (Suppl. Table S13). The other, non-DE, targets are similarly involved in signalling (*ephb1*), but also lipid metabolism (*acot11*, *cyp2j2*, *mylip*), and cytokinesis (*septin5*). Downregulated miRNAs targeted 10 mRNAs, that were therefore expressed at higher levels in HH offspring, of which the genes coding for immunoglobulin superfamily member 8 (*igsf8*), involved in immunity and cell adhesion, and Von Willebrand factor A domain containing 7 (*vwa7*) were also differentially expressed. For the non-DE targets, several functions in insulin signalling and energy metabolism (e.g. *atp6v0a1*, *irs2*, *ogdhl*), while others in vesicle-mediated transport and autophagy (e.g. *vps39*), crucial for removing damaged proteins and organelles that accumulate under heat stress, helping to restore cellular homeostasis. Therefore, unlike the widespread metabolic and stress-related changes induced by single-parent warming (HC/CH), biparental warming (HH) elicited a narrower but distinct miRNA-mRNA signature, prioritizing energy metabolism and signalling pathways (Fig. 5), implying non-additive parental effects on offspring post-transcriptional gene regulation.

**Figure 5.**
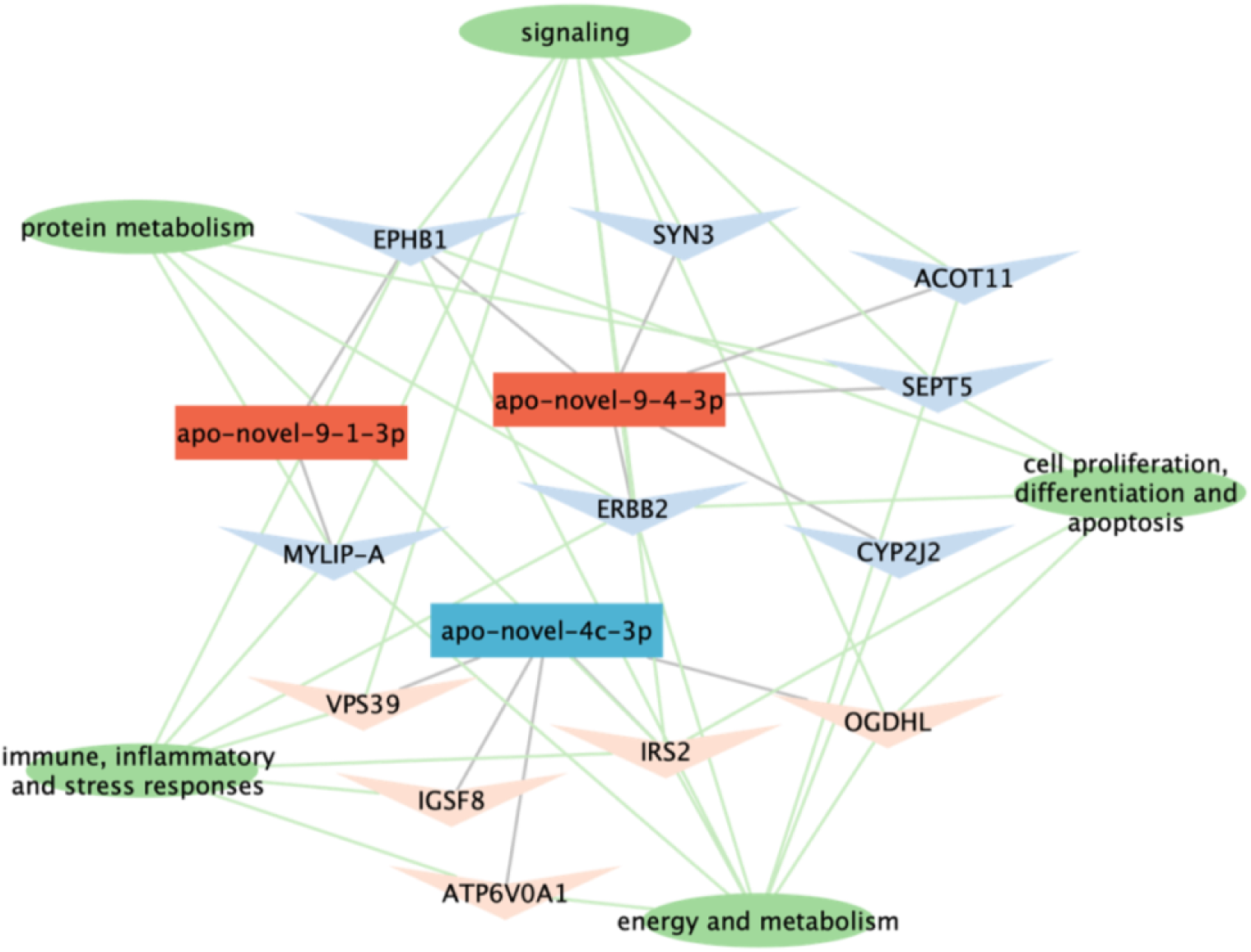
Regulatory network of DE-miRNAs, target genes and related functions in *A. polyacanthus* offspring from HH parents compared to offspring from CC parents. The quadrangular nodes represent miRNAs, the triangular nodes represent mRNAs, and the oval nodes represent functions and pathways. The red and blue nodes represent up- and downregulated RNAs in offspring from parents both raised at elevated temperature compared to offspring from control parents, respectively.

#### 3.2.2 Developmental effects of warming on miRNA expression

Offspring exposure to warming induced the differential expression of 23 miRNAs, regardless of their parents’ thermal history, revealing the direct role of developmental temperature in shaping miRNA-mediated gene regulation (Suppl. Table S5; Suppl. Fig. S5D). Among these, seven novel miRNAs were upregulated, but targeted only a total of six mRNAs. Apo-novel-1-1-5p expression in particular negatively correlated with two target mRNAs that were also differentially expressed: *itpr3* and *tef* (Suppl. Table S14). *Itpr3* encodes for inositol 1,4,5-trisphosphate receptor type 3, critical for calcium ion homeostasis and energy metabolism, and *tef*, a transcription factor, also showed reduced expression in offspring raised at warmer temperatures, indicating alterations in gene regulatory networks. Among the other, non-differentially expressed, targets of apo-novel-1-1-5p, *bco1* and *aldh1l1* are involved in vitamin and lipid metabolism, while *otud3*, targeted by the similarly upregulated apo-novel-34-5p, in energy metabolism. Conversely, the 16 downregulated miRNAs in warming-exposed offspring targeted 116 unique mRNAs (out of 122 significant DE miRNA-target mRNA pairs), with five of these also differentially expressed and upregulated. Among them, serpin family H member 1 (*serpinh1*), the only target of apo-miR-130c-3p, and integrin alpha-11 (*itga11*), targeted by apo-novel-15-3p, have roles in collagen binding and extracellular matrix organization. Additionally, Y-box binding protein 1 (*ybx1*), also targeted by apo-novel-15-3p, is a multifunctional DNA/RNA-binding protein, while AP-1 complex subunit mu01 (*ap1m1*), target of apo-novel-36-3p, functions in endocytosis and Golgi processing. The non-DE targets coded for proteins involved in immune system and inflammation (e.g. *ada2, ankib1, dusp4, ltn1, mgst1*), extracellular matrix remodelling and tissue repair (e.g. *col1a2, serpine2, yes1*), amino acid and lipid metabolism (e.g. *crot, degs1, dgat1, gls, gpt2, pla2g1*), transcription and gene expression (e.g. *hdac4, nr2f6, srsf11, tbl1xr1, ythdf2*), proteolysis (e.g. *fbxl19, lonp2, ubr4, zfand5*), protein folding in the ER (e.g. *retreg1, sdf2, tmtc3, tor1a*), cell cycle regulation and apoptosis (e.g. *birc6, cit, hebp1, rassf3, tnfrsf10b*). Hence, early-life exposure to elevated temperatures triggered a heterogeneous transcriptional regulation of a variety of key metabolic, quality control and tissue repair processes (Fig. 6).

**Figure 6.**
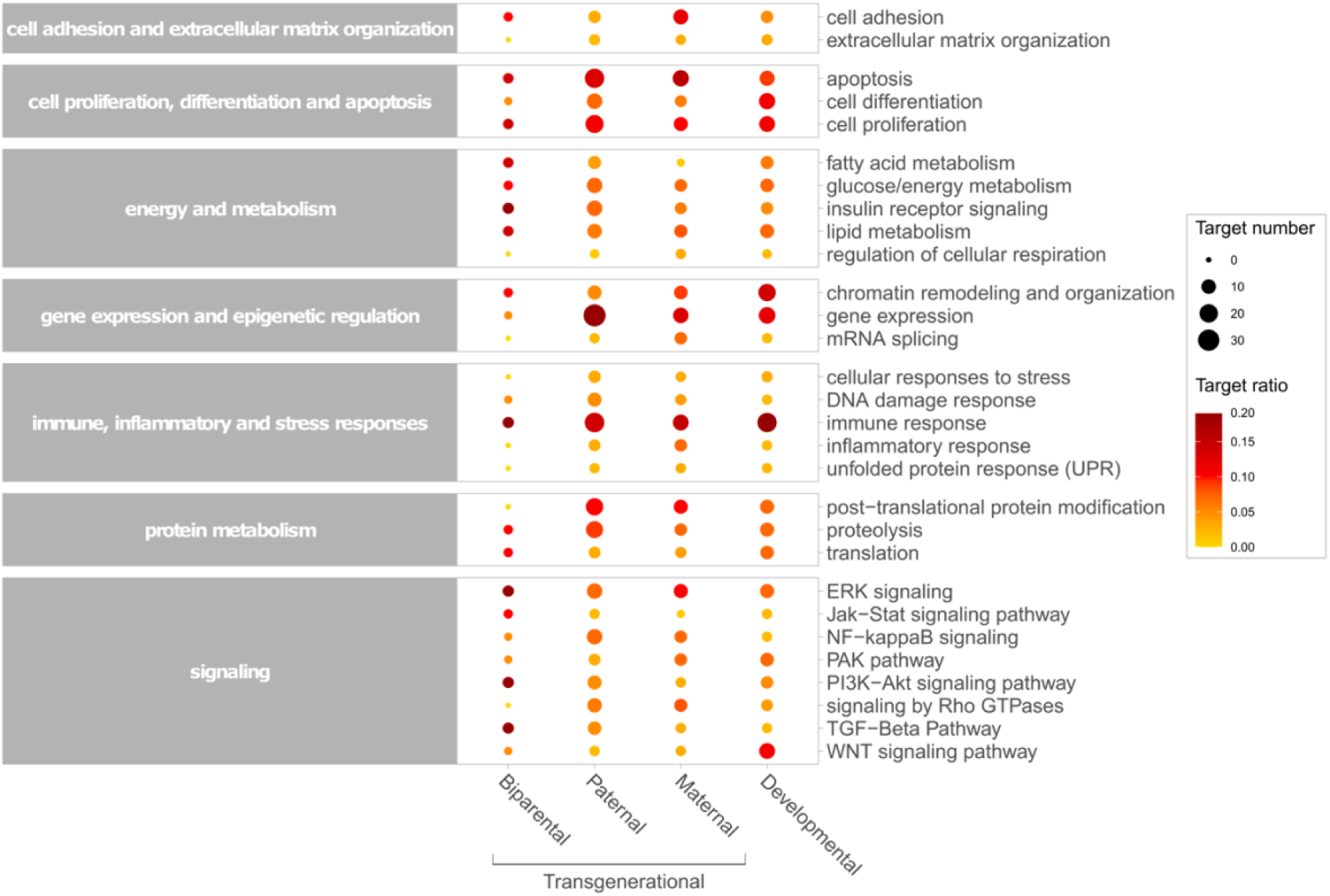
Functional representation of target transcripts of differentially expressed miRNAs due to the different transgenerational and developmental thermal exposures. The size and colour of the circles are proportional to the number and ratio of transcripts underlying each function, respectively.

## 4. DISCUSSION

Despite evidence that parental heat exposure influences offspring thermal tolerance, the epigenetic mechanisms and the relative contributions of maternal versus paternal effects are not well understood. Our study reveals that both developmental and transgenerational warming affect hepatic microRNA (miRNA) regulation in juvenile coral reef fish. While both exposures altered similar numbers of miRNAs, paternal warming alone - without concurrent maternal exposure - had the strongest inhibitory effects on gene expression, in addition to being the main driver of offspring gene expression (Bonzi et al., 2025). Most genes targeted by differentially expressed miRNAs in offspring from warming-exposed fathers were also significantly differentially expressed, indicating miRNAs strongly mediate paternal environmental programming. Maternal effects were present but less prominent. Importantly, combined paternal and maternal warming produced non-additive unique miRNA signatures, highlighting complex, non-linear interactions in transgenerational thermal responses.

Paternal warming caused the downregulation of conserved miRNAs such as apo-miR-375-3p and apo-miR-101a-3p in offspring, leading to increased expression of their target genes. MiR-375, a highly conserved regulator of metabolic processes across species, influences insulin secretion and glucose metabolism in mammals, and serves as a biomarker for metabolic diseases (Dumortier et al., 2020; Higuchi et al., 2015). Its expression responds to temperature, both in humans, where cold reduces miR-375 levels in adipose tissue regulating key thermogenic genes (Seeliger et al., 2022) and in fish, for example in largemouth bass *Micropterus salmoides*, where it has been linked to lipid metabolism modulation during heat stress (Zhao et al., 2024). Targets of apo-miR-375-3p found here such as secreted frizzled-related protein 3 (*sfrp3/frzb*) and thioredoxin related transmembrane protein 2 (*tmx2*) are involved in lipid metabolism (Guan et al., 2021) and mitochondrial/ER function. In fact, loss of *tmx2* impairs mitochondrial respiration, shifting energy metabolism under stress (Vandervore et al., 2019). *Tmx2* also interacts with calnexin to regulate protein folding and the unfolded protein response (UPR) to prevent ER stress due to an accumulation of misfolded proteins (Hetz et al., 2020, Vandervore et al., 2019). Therefore, miR-375 role as thermal-responsive regulator is maintained also in *A. polyacanthus*, where it mediates transgenerational metabolic reprogramming in response to paternal warming via mitochondrial and ER homeostasis. Similarly, the conserved metabolic regulator mir-101 (Higuchi et al., 2015), known to modulate mitochondrial activity (Ziemann et al., 2022) and to respond to environmental stress (Sun et al., 2017; Yan et al., 2024), here caused the upregulation of genes (e.g. *calr, cnpy2, edem1*) involved in the UPR, as well as in tissue repair, indicating roles in stress adaptation and regenerative processes. Additionally, mir-101 targeted lipid-energy metabolism genes (e.g. *acsl6, prkab1*) in *A. polyacanthus* offspring, with its downregulation therefore suggesting increased metabolic activity to meet elevated energy demands. Overall, these miRNAs thus act as conserved thermal regulators, mediating transgenerational metabolic and stress responses to warming. Indeed, while showing enhanced metabolic flexibility, offspring from warming-exposed fathers may also be experiencing latent stress, evidenced by upregulated stress pathways and reduced growth, suggesting a trade-off between resilience and costs of transgenerational plasticity (Bonzi et al., 2025; Spinks et al., 2022). These miRNAs thus serve as key mediators linking environmental cues to phenotypic adjustments, balancing adaptive acclimation and underlying stress in a warming climate.

Compared to the paternal effects, maternal-only warming exposure caused more miRNAs to be differentially expressed in the offspring, though these targeted fewer genes. MiR-1-3p, a known biomarker for heat stress in fish (Lin & Meegaskumbura, 2025) with functions in cell growth, immune response and glucose metabolic regulation (Ma et al., 2019; Qiang et al., 2017; Sun et al., 2019; Zhou et al., 2019), was upregulated in offspring from heat-exposed mothers and control fathers. In *A. polyacanthus*, mir-1-3p targeted genes associated with immune response (e.g. *inpp5d*), tissue organization (e.g. *ctnnal1, rara*), and energy metabolism (e.g. *ehd1, galm, hmga1*), thus downregulated in the offspring, suggesting its role extends beyond acute stress response to transgenerational thermal acclimation. Similarly, miR-27b-1-3p, also upregulated, targeted genes involved in energy homeostasis (e.g. *nubpl*) and immune function (e.g. *havcr1*, *tmed1*). Its involvement in fatty acid metabolism, stress and immune signaling pathways (observed in zebrafish and mammals; van Gelderen et al., 2022; Zheng et al., 2014) indicates a conserved role in thermal stress response, though its dysregulation has also been linked to liver disease, raising concerns about potential trade-off costs (Cheng et al., 2023; López-Pastor et al., 2021; Zhao et al., 2023). Accordingly, a novel miRNA upregulated because of maternal exposure to warming, apo-novel-37-3p, also targeted genes important in tissue homeostasis and wound healing (e.g. *ddr1*, *itga5*), as well as others key to the activation of inflammatory and adaptive immune responses (e.g. *card8*, *card11*). Among the downregulated miRNAs due to maternal warming, on the other hand, we found miRNAs mostly lacking significant targets in *A. polyacanthus*, implying either functional divergence or compensatory regulatory mechanisms. An exception was apo-miR-19da-3p, which targeted genes related to apoptosis (e.g. *nf2*, *pcdh17*), intracellular transport and cytoskeleton modifications (e.g. *agap1, iqsec1*), as well as gene expression and post-transcriptional processes (e.g. *ankrd11*, *celf1, ddx17*). Overall, these maternal molecular signatures suggest an overall suppression of costly processes (e.g. immunity, tissue repair) to prevent excessive inflammation under heat stress, accompanied by shifts in energy allocation away from tissue maintenance toward stress tolerance mechanisms, like apoptosis (Duan et al., 2024; Liu et al., 2022) and gene expression regulation. While beneficial for heat tolerance, these changes may reduce immune competence and metabolic flexibility, likely impacting long-term susceptibility to secondary stressors (e.g. pathogens, injuries), which might hinder the ability of new generations to survive long-term in a warming ocean (Aversa-Marnai et al., 2025; Scharsack & Franke, 2022).

The trade-offs between acclimatory benefits and physiological costs were especially pronounced in offspring from thermally mismatched parents (i.e., only one parent exposed to warming) compared to those with both parents acclimated to elevated temperatures, a pattern driven by the miRNA expression shifts linked to parental thermal history interactions. In particular, mismatches between parental thermal experiences activated miRNA-mRNA pathways linked to hepatic diseases and stress responses. Downregulated miRNAs in offspring from mismatched parents led to the overexpression of genes involved in apoptosis and tissue repair (e.g. *ddx17, nf2, notch2, rbm25*), suggesting hepatic damage. Moreover, these miRNAs also targeted genes involved in inflammation and stress, such as ARF GTPase activating protein 1 (*arfgap1*), a component of the cellular response to stress and the UPR, TAO kinase 1 (*taok1*), which regulates DNA damage responses (Raman et al., 2007) and is a marker for non-alcoholic fatty liver disease in humans (Xia et al., 2023), as well as the stress-activated TRAF3 interacting protein 2 (*traf3ip2*), which mediates inflammatory signals leading to tissue injury and dysfunction (Erikson et al., 2017; Wei et al., 2022). On the contrary, offspring from parents that were both exposed to warming showed an overall activation of adaptive miRNA-mediated regulation, with the differential expression of a unique set of miRNAs, novel to *A. polyacanthus*. For example, among the targeted genes by the upregulated apo-novel-9-4-3p and, consequently, suppressed in these offspring, was acyl-CoA thioesterase 11 (*acot11*). This metabolic regulator is crucial in energy homeostasis (Desai et al., 2018; Zhang et al., 2012), and exhibits temperature-dependent expression in mice, increasing during cold exposure and decreasing with warm acclimation (Adams et al., 2001). Interestingly, while chronic stress typically upregulates *acot11* to promote energy conservation (Xu et al., 2024), this response can become maladaptive over time. Supporting this, mice lacking *acot11* show increased energy expenditure and reduced ER stress responses, and are protected against metabolic disorders like insulin resistance and non-alcoholic fatty liver disease (Zhang et al., 2012). The observed miRNA-mediated *acot11* suppression in *A. polyacanthus* offspring from parents that were both exposed to warming may therefore represent an adaptive mechanism to counteract detrimental metabolic effects of chronic thermal stress. Another potential adaptive acclimation of biparentally exposed offspring to future ocean warming conditions involves apo-novel-9-1-3p, an upregulated miRNA targeting the myosin regulatory light chain interacting protein (*mylip*). Notably, *mylip* suppression in fish enhances hypoxia tolerance by upregulation of the hypoxia signalling pathway and the inhibition of the proteasomal degradation of hypoxia-inducible factors (Li et al., 2025), which may help offspring of warming-exposed parents cope with ocean deoxygenation caused by climate change (Keeling et al., 2010; Pezner et al., 2023). Together, these findings highlight how parental thermal histories shape offspring miRNA expression to fine-tune metabolic, immune, and stress-related pathways, enabling adaptive responses when parental environments match, but potentially predisposing offspring to hepatic dysfunction and metabolic stress when parental thermal experiences conflict. Notably, the results of the miRNA analysis are consistent with both our previous findings of maladaptive responses in offspring from mismatched parents on overall gene expression (Bonzi et al., 2025) and physiological measurements (Spinks, 2021), and with the theoretical expectation of beneficial acclimation when the same environmental cue is perceived multiple times, so that it can overcome potential detection limits (Bell & Hellmann, 2019). Indeed, biparental miRNA-mediated regulation may help offspring offset the energetic costs of warming, indicating that matching parental exposure can equip the next generation with transcriptional tools to cope with increased energy demands and mitigate the negative effects of warmer water.

Developmental exposure to warming induced the most extensive miRNA differential regulation, targeting the largest subset of mRNAs. However, only a few of the targeted transcripts were also significantly differentially expressed, suggesting a more subtle regulatory role of miRNAs in developmental thermal plasticity compared to both single parental effects, particularly the paternal ones. Among the differentially expressed targets were several genes involved in energy metabolism, indicating metabolism adjustment to meet the increased energetic demand at higher temperature, but also serpin family H member 1 (*serpinh1* or heat shock protein 47), an ER-resident chaperone critical for collagen biosynthesis and a known heat-stress marker in fish (Akbarzadeh et al., 2018; Ignatz et al., 2024; Wang et al., 2016). Its upregulation under heat shock is part of the cellular response to unfolded proteins and, in mammals, *serpinh1* overexpression is linked to liver fibrosis, where excessive collagen deposition occurs due to chronic injury (Ito & Nagata, 2017). Similarly, both acute and prolonged heat exposure in fish has been associated with liver damage (Duan et al., 2024; Han et al., 2023; Yan et al., 2025), and developmental warming in clownfish (*Amphiprion ocellaris*) triggers fibrotic gene activation (Moore et al., 2024). Consistent with this, we observed miRNA-mediated upregulation of fibrosis-related genes in offspring developmentally exposed to warming, such as *col1a2* (collagen production), *ada2* (triggers PDGF-B release, activating collagen production; Tiwari-Heckler et al., 2021), *itga11* (upregulated in liver injury, causes excess collagen production; Bansal et al., 2017), and *ybx1* (increases extracellular matrix deposition and enhances pro-fibrotic genes chromatin accessibility; Tang et al., 2023). Therefore, direct experience of elevated water temperature, regardless of the parental thermal history, caused the activation of miRNA-mRNA regulatory networks that are not only involved in expected metabolism adjustment to overcome the increased energy requirements, but that could also represent detrimental effects for the offspring’s fitness, particularly fibrotic liver damage, which might be further exacerbated when the parental thermal histories do not coincide.

Epigenetic-mediated transgenerational plasticity could represent a fast acclimation strategy to anthropogenic climate change. Here, we show that in the coral reef fish *A. polyacanthus* both parents are able to elicit differential expression of miRNAs in their offspring following warming exposure, with some transcriptional responses potentially adaptive for the offspring in the warmer environment, and paternal exposure eliciting stronger regulatory changes, in agreement with previous studies on miRNA-mediated paternal effects (Gapp et al., 2014; Rodgers et al., 2013, 2015; Wang et al., 2021; Yin et al., 2025). However, parental effects were not additive and were unique to each parental combination, emphasizing the critical need to consider maternal, paternal, and interactive effects when assessing the potential for epigenetic-mediated acclimation to environmental change. Since limited to juvenile liver tissue, future studies should explore whole-organism responses across life stages and experimentally validate miRNA roles (e.g. via knockdown) to confirm the adaptive significance of these findings. Nevertheless, our work highlights miRNAs as mediators of climate resilience, and underscores the complex trade-offs of transgenerational plasticity when parental environments mismatch, a critical consideration for predicting species’ adaptive capacity under rapid environmental change.

## DATA ACCESSIBILITY

Both RNA-seq mRNA and miRNA data can be found under the BioProject PRJNA998209. Electronic supplementary material is available online.

## ETHICS

All procedures were performed in accordance with relevant guidelines and were conducted under James Cook University’s animal ethics approval A1990, A2210 and A2315.

## AUTHOR CONTRIBUTIONS

L.C.B., J.M.D. and R.K.S. conceptualized the experiment and collected the samples. Fish rearing was overseen by L.C.B. and R.K.S. L.C.B. prepared the samples for sequencing, performed the data analysis and drafted the initial version of the manuscript with input from C.S. Funding was obtained by C.S., J.M.D., R.K.S., P.L.M. and T.R. All authors reviewed, provided feedback, and approved the final version for publication.

## ACKNOWLEDGEMENTS

We thank the Marine and Aquaculture Research Facilities Unit at JCU and KAUST Bioscience Core Lab, as well as all members of Prof Schunter’s Lab at HKU for their feedback and support.

