## Supplementary Figures for "MicroRNA-mediated transgenerational plasticity reveals pathways for ocean warming resilience"

^2^ ARC Centre of Excellence for Coral Reef Studies, James Cook University, Australia

^4^ Blue Carbon Section, Australian Government Department of Climate Change, Energy, the Environment and Water, Australia

**Supplementary Figures**

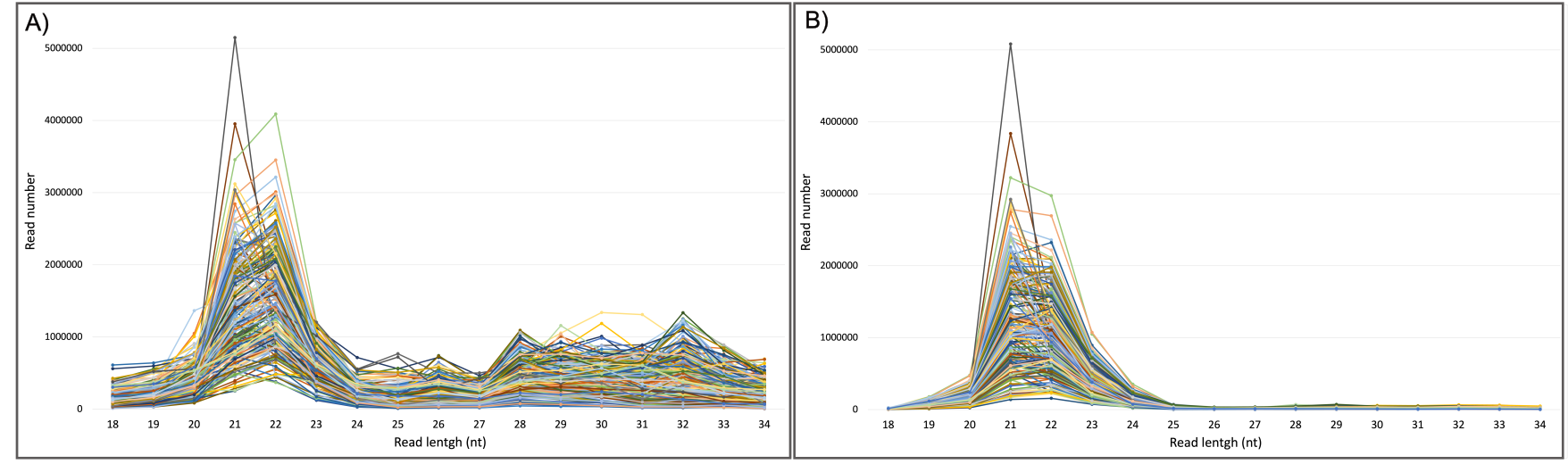

**Supplementary Figure S1.** Length distribution of sequenced *Acanthochromis polyacanthus* miRNAs, A) before and B) after non-miRNA small RNA filtering. Each colored line represents a different sample.

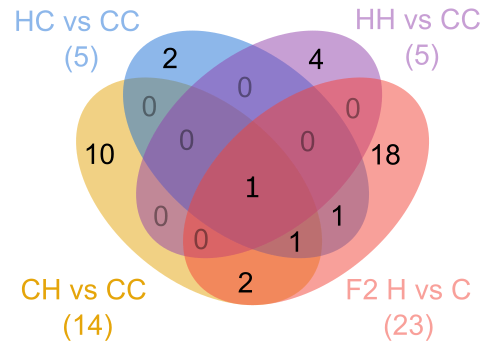

**Supplementary Figure S2.** Venn diagram of the miRNA differential expression (DE) in pairwise comparisons between offspring raised at elevated vs control temperature (F2 H vs C) and between offspring from different parental thermal combinations. The first letter stands for the paternal thermal environment, the second for the maternal one; ‘C’ = control temperature and ‘H’ = +1.5°C.

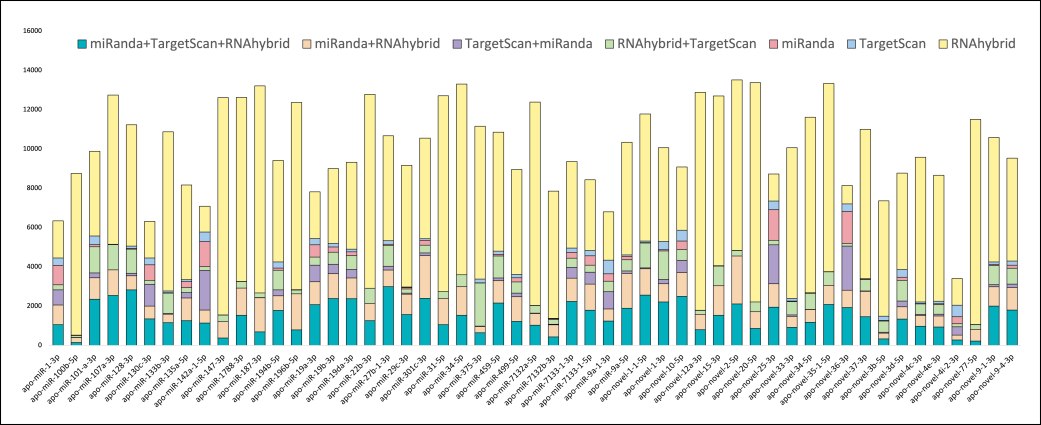

**Supplementary Figure S3.** Predicted putative target mRNAs for each differentially expressed miRNA by three in silico tools, TargetScan, miRanda, and RNAhybrid.

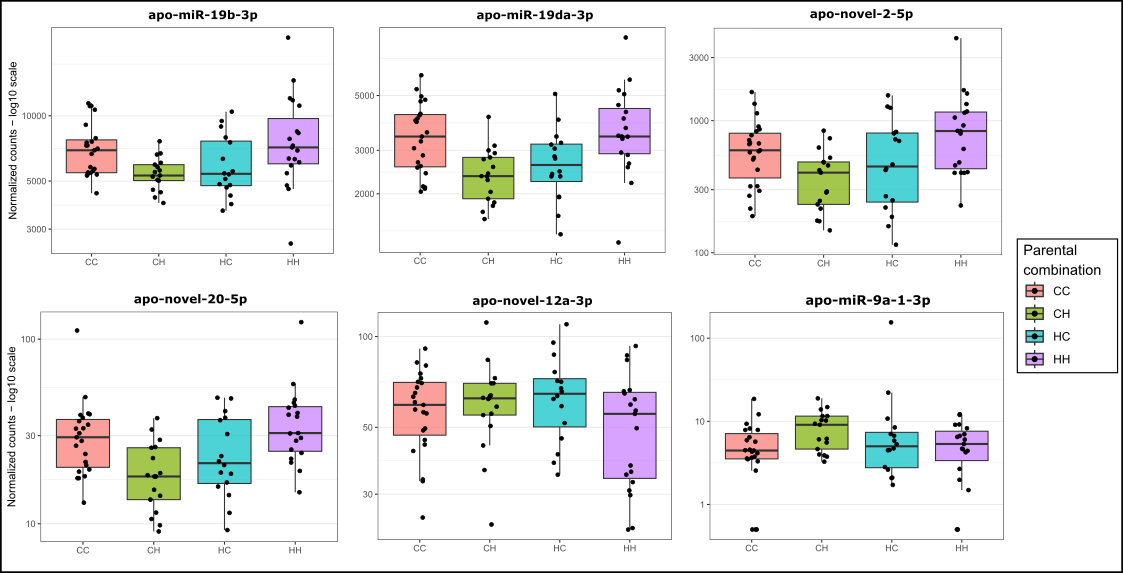

**Supplementary Figure S4.** Expression profiles of differentially expressed miRNAs showing interactive effects between maternal and paternal thermal exposures. In the parental combinations, the first letter stands for the paternal thermal environment, the second for the maternal one; ‘C’ = control temperature and ‘H’ = +1.5°C.

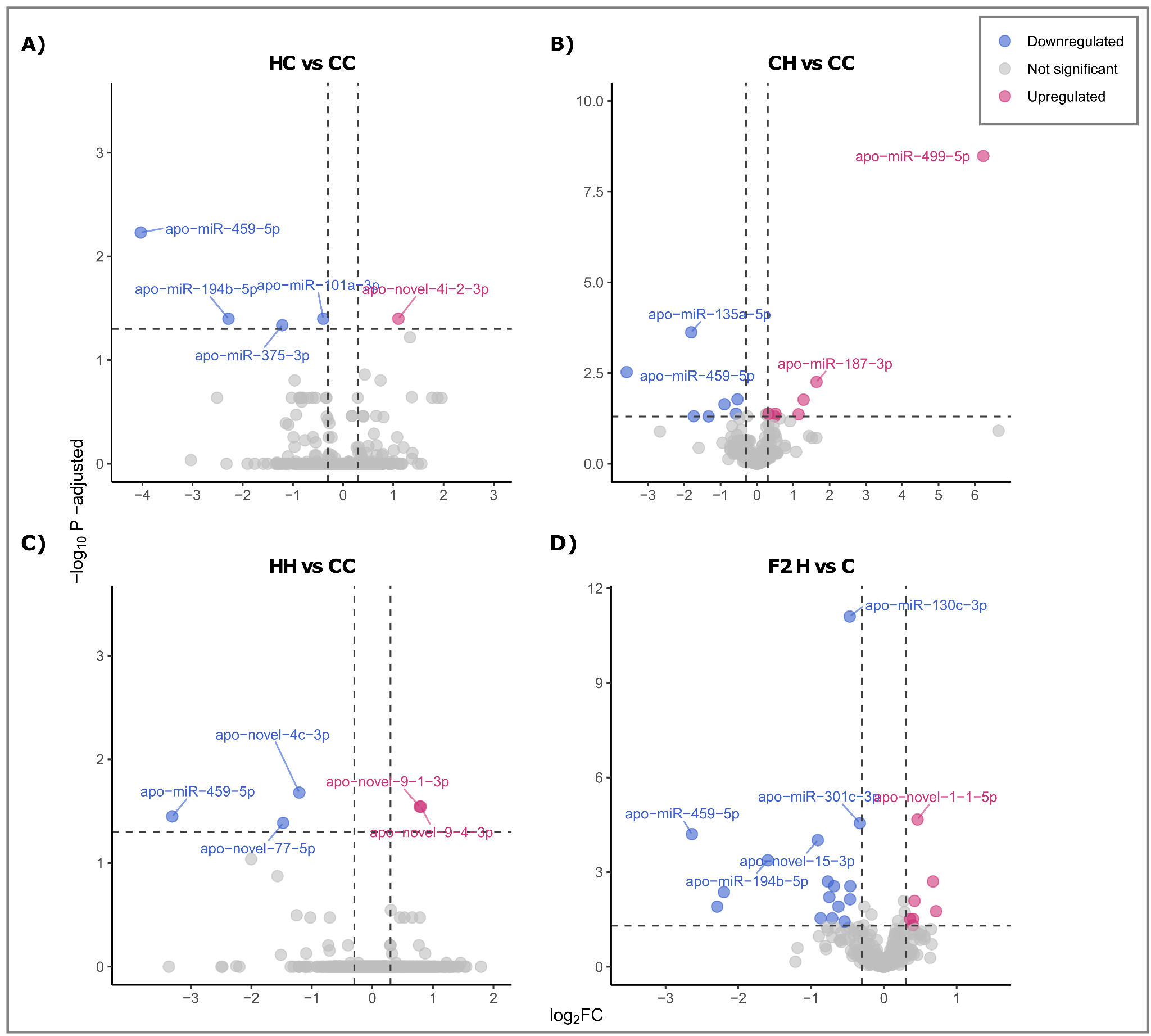

**Supplementary Figure S5.** Volcano plots of differentially expressed miRNAs in offspring A) from fathers exposed to warming and mothers to control (HC vs CC), B) from fathers exposed to control and mothers to warming (CH vs CC), C) from parents both exposed to warming (HH vs CC), regardless of their own developmental thermal regime and D) developmentally exposed to warming, regardless of their parental thermal history (F2 H vs C).
